# Retinal input is required for the maintenance of neuronal laminae in the ventral lateral geniculate nucleus

**DOI:** 10.1101/2024.01.12.575402

**Authors:** Katelyn Stebbins, Rachana Deven Somaiya, Ubadah Sabbagh, Yanping Liang, Jianmin Su, Michael A. Fox

**Affiliations:** Fralin Biomedical Research Institute at Virginia Tech Carilion, Roanoke, Virginia, 24016, USA; Graduate Program in Translational Biology, Medicine, and Health, Virginia Tech, Blacksburg, Virginia, 24061, USA; Virginia Tech Carilion School of Medicine, Roanoke, 24016, USA; Department of Molecular and Cell Biology, University of California Berkeley, Berkeley, California, 94720, USA; McGovern Institute for Brain Research, Department of Brain and Cognitive Sciences, Massachusetts Institute of Technology, Cambridge, MA, USA; School of Neuroscience, College of Science, Virginia Tech, Blacksburg, Virginia, 24061, USA; Department of Biological Sciences, College of Science, Virginia Tech, Blacksburg, Virginia, 24061, USA; Department of Pediatrics, Virginia Tech Carilion School of Medicine, Roanoke, Virginia, 24016, USA; Department of Biology, College of Natural Sciences, University of Massachusetts Amherst, Amherst, Massachusetts, 01003, USA

**Keywords:** lateral geniculate nucleus, retinal ganglion cells, GABAergic neuron, retina, visual thalamus

## Abstract

Retinal ganglion cell (RGC) axons provide direct input into several nuclei of the mouse visual thalamus, including the dorsal lateral geniculate nucleus (dLGN), which is important for classical image-forming vision, and the ventral lateral geniculate nucleus (vLGN), which is associated with non-image-forming vision. Through both activity- and morphogen-dependent mechanisms, retinal inputs play important roles in the development of dLGN, including the refinement of retinal projections, morphological development of thalamocortical relay cells (TRCs), the timing of corticogeniculate innervation, and the recruitment of inhibitory interneurons from progenitor zones. In contrast, little is known about the role of retinal inputs in the development of vLGN. Grossly, vLGN is divided into two domains, the retinorecipient external vLGN (vLGNe) and the non-retinorecipient internal vLGN (vLGNi). We previously found that vLGNe consists of transcriptionally distinct GABAergic subtypes that are distributed into at least four adjacent laminae. At present, it remains unclear whether retinal inputs influence the development of these cell-specific neuronal laminae in vLGNe. Here, we elucidated the developmental timeline for the formation and maintenance of these laminae in the mouse vLGNe and results indicate that these laminae are specified at or before birth, well before eye-opening and the emergence of experience-dependent visual activity. We observed that mutant mice without retinal inputs have a normal laminar distribution of GABAergic cells at birth; however, after the first week of postnatal development, these mutants exhibited a dramatic disruption in the laminar organization of inhibitory neurons and clear boundaries between vLGNe and vLGNi. Overall, our results show that while the formation of cell type-specific layers in vLGNe does not depend on RGC inputs, retinal signals are critical for their maintenance.

## INTRODUCTION

In the vertebrate visual system, axons of retinal ganglion cells (RGCs) convey visual information from the world to numerous distinct regions of the hypothalamus, thalamus, and midbrain (Monavarfeshani et al., 2017; Morin and Studholme, 2014; Muscat et al., 2003). In rodents, RGC axons innervate a region of the visual thalamus known as the lateral geniculate complex, which is composed of three nuclei: the dorsal lateral geniculate nucleus (dLGN), the ventral lateral geniculate nucleus (vLGN), and the intergeniculate leaflet (IGL) (Fleming et al., 2006; Gaillard et al., 2013; Martersteck et al., 2017; Monavarfeshani et al., 2017; Morin and Studholme, 2014). In recent years, anatomical and functional studies have unmasked the complexity of the role of visual thalamus in the processing of visual information, although the bulk of these efforts have focused on dLGN, which connects the retina to the visual cortex and is associated with image-forming vision (Guido, 2018; Tiriac et al., 2018; Huberman et al., 2008).

In dLGN, numerous studies have revealed details about the developmental timeline of thalamocortical relay cells, the migration of local interneurons, the arrival of other types of axons, and the emergence and impact of experience-dependent visual activity on each of these processes. During perinatal mouse development, RGC axons establish connections with the dLGN, with axons from the contralateral eye reaching the dLGN between embryonic day 16 and birth (E16-P0), preceding the arrival of axons from the ipsilateral eye (P0-P2) (Godement et al., 1980). By the time eye opening occurs, approximately around postnatal day 12 to 14 (∼P12-P14), the eye-specific domains become well-segregated and resemble the mature pattern of innervation seen in adults (Huberman et al., 2008). The refinement of these eye-specific domains and the formation of retinotopic maps involves the remodeling of individual axon arbors, where axons projecting to inappropriate target areas are retracted or eliminated, while those projecting to the correct locations are matured and maintained (Dhande et al., 2011). The mechanisms underlying retinotopic map formation and eye-specific axon segregation have been well studied and include complex interactions between developmentally relevant molecular cues and both spontaneous and experience-dependent neural activity (Cang et al., 2008; Pfeiffenberger et al., 2005, 2006; Huberman et al., 2008).

For many years, it has been appreciated that retinal inputs not only convey neuronal signals from the retina to the dLGN, but also influence thalamic development. Once RGC axons arrive into visual thalamus, they contribute critical roles in shaping the development of several distinct cell types and non-retinal circuits in dLGN. For example, retinal inputs play a crucial role in the morphological development of principal neurons within dLGN, and their loss disrupts the dendritic growth of these neurons (Cheadle et al., 2018; Chen and Regehr, 2000; El-Danaf et al., 2015). Perhaps more surprisingly, retinal inputs are also critical for the recruitment of local GABAergic interneurons from thalamic and tectal progenitor zones in dLGN (Charalambakis et al., 2019; Golding et al., 2014; Jager et al., 2016, Somaiya et al., 2022). Moreover, retinal inputs influence the timing of the arrival of other types of non-retinal axons, including corticogeniculate axons, which are the most abundant axons that innervate thalamocortical relay cells in dLGN (Brooks et al., 2013; Seabrook et al., 2013).

In stark contrast to this, very little is known about the development of cell types and associated circuitry in vLGN. This knowledge gap has been due, in part, to the paucity of what is known about vLGN cytoarchitecture and neuronal cell types. One recent study accounted for at least six distinct types of GABAergic neurons in vLGN, many of which organize into hidden subtype-specific laminae and appear to receive direct retinal inputs (Sabbagh et al., 2021). Here, we sought to document the development of these vLGN laminae and based on the critical role of retinal inputs in orchestrating aspects of dLGN development, we asked whether retinal inputs are required for establishing laminae within the retinorecipient vLGN. Surprisingly, we discovered that retinal inputs are not required for the initial formation of vLGN laminae, but instead are required for their maintenance.

## MATERIALS AND METHODS

### Animals

Wild-type C57BL/6 mice were obtained from Jackson Laboratory. *Math5*^−/−^ mice (stock #042298-UCD) (Wang et al., 2001) were obtained from S. W. Wang (University of Texas MD Anderson Cancer Center), and *Fgf15*^−/−^ (stock #032840-UCD) were obtained from the Jackson Laboratory. Animals were housed in a temperature–controlled environment, in a 12 hr dark/light cycle, and with access to food and water ad libitum. Both male and female mice were used in the experiments. All animal procedures were performed in accordance with the university’s animal care committee’s regulations. Unless otherwise stated, n = number of animals, and a minimum of three age- and genotype-matched animals were compared in all experiments.

### Riboprobe production

Riboprobes against *Lypd1* (Lypd1-F: AAGGGAGTCTTTTTGTTCCCTC; Lypd1-R: TACAACGTGTCCTCTCAGCAGT; NM_145100.4; nucleotides 849-1522, NM_007492.4; nucleotides 1830-2729), *Ecel1* (Ecel1-F: CGCGCTCTTCTCGCTTAC; Ecel1-R: GGAGGAGCCACGAGGATT; NM_001277925.1; nucleotides 942-1895), and *Penk* (Penk-F: TTCCTGAGGCTTTGCACC; Penk-R: TCACTGCTGGAAAAGGGC; NM_001002927.3; nucleotides 312-1111) were generated in-house. cDNAs were generated using Superscript II Reverse Transcriptase First Strand cDNA Synthesis kit (#18064014, Invitrogen) according to the manufacturer manual, amplified by PCR using primers designed for the above gene fragments, gel purified, and then cloned into a pGEM-T Easy Vector using pGEMT Easy Vector kit, (#A1360, Promega) according to the kit manual. Antisense riboprobes against target genes were synthesized from 5 μg linearized plasmids using digoxigenin-(DIG) or fluorescein-labeled uridylyltransferase (UTP) (#11685619910, #11277073910, Roche, Mannheim, Germany) and the MAXIscript in vitro Transcription Kit (#AM1312, Ambion), according to the manufacturer’s instructions. Five micrograms of riboprobe (20 μl) were hydrolyzed into ∼0.5 kb fragments by adding 80 μl of water, 4 μl of NaHCO3 (1 M), and 6 μl Na2CO3 (1 M), and incubating the mixture at 60°C. The RNA fragments were precipitated in ethanol and resuspended in RNAase-free water.

### *In-situ* hybridization (ISH)

ISH was performed on 16μm thin cryosections. Sections were first allowed to air dry for 1hr at room temperature and washed with PBS for 5 min to remove freezing media. They were then fixed in 4% PFA for 10 min, washed with PBS for 15 min, incubated in proteinase K solution for 10 min, washed with PBS for 5 min, incubated in 4% PFA for 5 min, washed with PBS for 15 min, incubated in acetylation solution for 10 min, washed with PBS for 10 min, incubated in PBS-diluted 0.1% triton for 30 min, washed with PBS for 40 min, incubated in 0.3% H_2_O_2_ for 30 min, washed with PBS for 10 min, pre-hybridized with hybridization solution for 1 hr, and then hybridized with heat-denatured riboprobes at 62.5°C overnight. The sections were then washed five times in 0.2X SSC buffer at 65°C. Slides were then washed with TBS, blocked, and incubated with horseradish peroxidase-conjugated anti-DIG (#11426346910, Roche) or anti-fluorescent antibodies (#11207733910, Roche) overnight at 4°C. Finally, the bound riboprobes were detected using a tyramide signal amplification system (#NEL753001KT, PerkinElmer).

### Intraocular injection of CTB

We anesthetized mice with isoflurane or hypothermia for intraocular injections. A fine glass pipette attached to a picospritzer was used to intravitreally inject 1–2 μl of CTB (1 mg/ml) into the eye. After 2-3 days, the mice were perfused, and their PFA-fixed brains were sectioned (100 μm) using a vibratome (HM650v, Thermo Fisher Scientific), stained with 4′,6-diamidino-2-phenylindole (DAPI) (1:5,000 in water), and mounted using Vectashield (Vector Laboratories, RRID:AB_2336789).

## QUANTIFICATION AND IMAGING

### Imaging

All imaging for quantification was performed on a confocal Zeiss LSM 700 microscope at 20x magnification and 0.5 digital zoom. Each representative image in the figure is a maximum-intensity projection. Colocalization was confirmed using single-plane images. Cell counts were obtained from three sections within the middle of each region. When comparing sections from different age groups or genotypes, images were acquired with identical parameters. A minimum of three animals (per genotype and per age) were compared in all experiments.

### Area of vLGNe and vLGNi

To measure the area of vLGNe and vLGNi in wild-type and *Math5*^-/-^ animals, the entire vLGN was traced with ImageJ software (version 1.52n, NIH), and the total area was measured. For this purpose, the lateral boundary of the vLGN and of the vLGNe was defined by the optic tract and the change in density of DAPI^+^ cells were used to discern the medial border of vLGNi (Figure 1A, arrows) since previously identified markers that exclusively label neurons in vLGNi do not reliably label cells in the neonatal brain (data not shown). The medial border of vLGNe was identified by discerning the boundary of CTB localization. The area of vLGNi was determined by subtracting the previously determined area of vLGNe from the overall area of vLGN.

**Figure 1.**
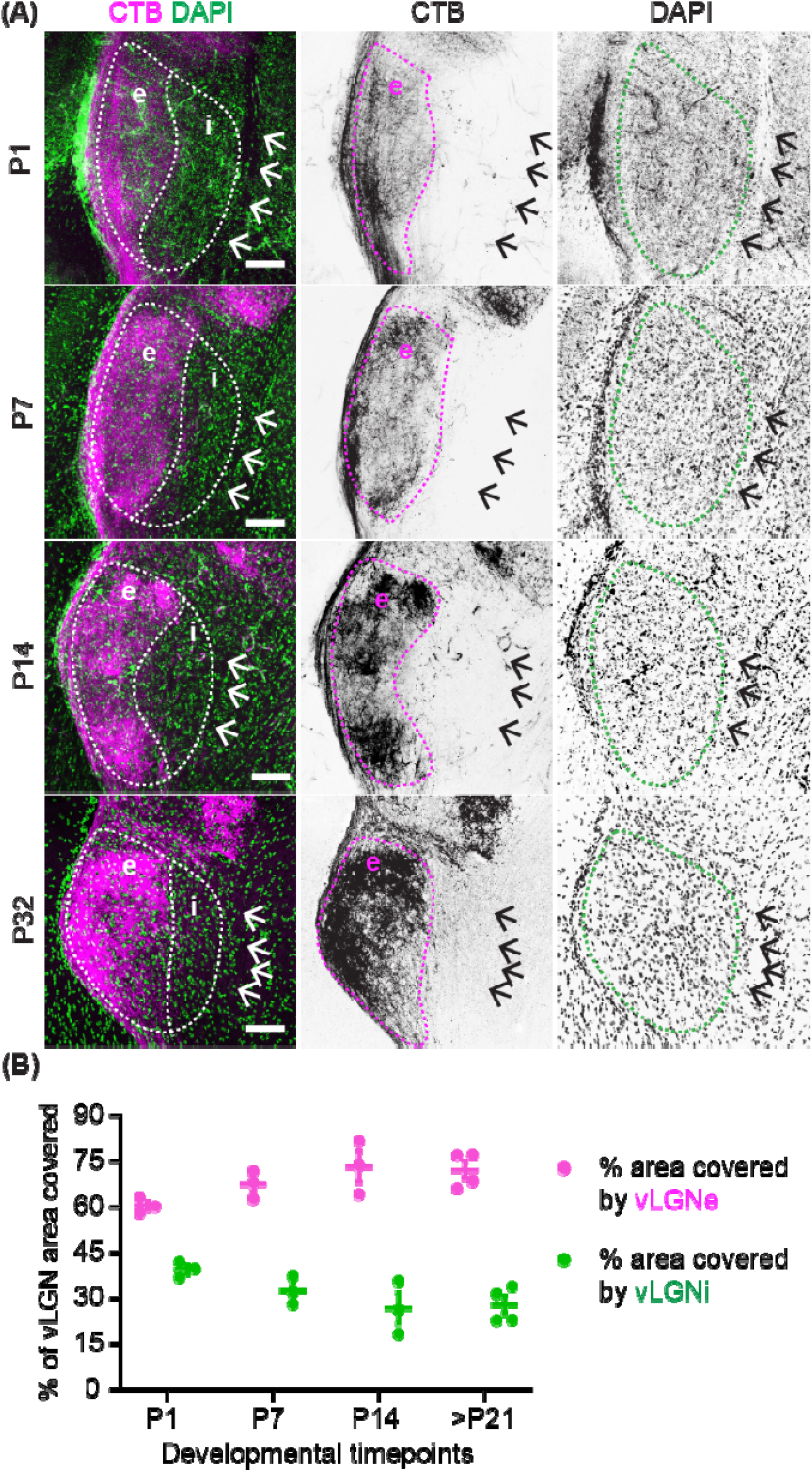
Retinal input is necessary for the developmental expansion of vLGNe. (A) Retinal terminals in vLGNe labeled by monocular injection of fluorescently-conjugated CTB at P1, P7, P14, and P32 of wild-type (WT) thalamus. Medial border of vLGNi was determined by cytoarchitecture (see black arrows in rightmost panels). (B) Quantification of the percentage of vLGN area covered by vLGNe and vLGNi across developmental time points in WT mice. Each data point represents one biological replicate and bars depict means ± SEM (P1, P7, P14 n=3; >P21 n=4). Scale bars in A: 100 μm.

### Quantification of laminar organization

To quantify the spatial distribution of cell-type marker expression across the entire vLGN, we used line scan script that runs in ImageJ and overlays the vLGN with equally spaced lines. A summary of how this script works: confocal images are first thresholded and binarized to determine the curvature of the vLGN in a particular image, the script prompts the user to draw a line along the optic tract adjacent to the vLGN, then automatically draws lines of a set length and number guided by that curve and plots detected signal across the x-coordinates of each line. Signal values at each x-coordinate are then averaged to determine where there is specific enrichment for that marker in that spatial zone of the vLGN.

### Percentage of *Ecel1*^*+*^, *Lypd1*^*+*^, and *Penk*^*+*^ cells

Images were obtained using 20x magnification and 0.5 digital zoom. We quantified *Ecel1*^+^, *Lypd1*^+^, and *Penk*^+^ (by ISH) cells and divided by the total number of DAPI^+^ cells counted in that tissue section. The “Count Tool” function in Adobe Photoshop (version: 21.1.2) was utilized for counting purposes.

## RESULTS

### Development of the external and internal divisions of vLGN

The rodent vLGN can be divided into two distinct domains based on cytoarchitecture: the retinorecipient external division (vLGNe) and the non-retinorecipient internal division (vLGNi). A recent study identified and characterized novel GABAergic neuronal types that occupy these regions (Sabbagh et al., 2021). To investigate the development of the cell types in the neo- and perinatal vLGN, we assessed cytoarchitectural changes in these vLGN subdivisions during the first three weeks of mouse development. We identified these regions based on DAPI labeling and anterograde labeling of retinal axons. Retinal terminals were labeled by intraocular injection of fluorescently conjugated CTB, a tracer that efficiently labels retinal projections into the brain (Muscat et al., 2003). The area occupied by retinal axons was determined to be vLGNe (Sabbagh et al., 2021; Govindaiah et al., 2023; Guido, 2018). We observed that the proportion of vLGN represented by vLGNe increased with postnatal age, while the percentage of vLGN area represented by vLGNi decreased (Figure 1A-B). At P1, the proportions of area occupied by vLGNe and vLGNi were 60% and 40%, respectively. After P21, the difference between the proportions of vLGNe and vLGNi doubled, with vLGNe occupying 72% of vLGN and vLGNi occupying only 28% of vLGN. This suggests that the arrival of retinal inputs potentially influences the expansion of vLGNe during development.

### Subtype-specific laminae are specified in the neonatal vLGN

We next set out to characterize the development of neuronal cell types in vLGN, as well as explore when cell-type specific laminae form in vLGN. A previous study characterized several vLGN cell types based on the generation of specific transcripts. vLGNe can be further divided into at least four subtype-specific layers, each composed of molecularly and functionally distinct neurons, several of which receive direct monosynaptic input from the retina (Sabbagh et al., 2021). In some cases, the expression of these transcripts was shown to be developmentally regulated, making it potentially challenging to use riboprobes against those genes to study laminar organization at different developmental ages. This was the case for both *Ecel1*^*+*^ and *Lypd1*^*+*^ GABAergic neurons, which are present in the same region (Figure 2A-B) within the vLGN but exhibited different developmental expression patterns, with *Lypd1* expression decreasing with development and *Ecel1* expression being low or not detectable at early postnatal ages (Sabbagh et al., 2021). To determine if these markers were generated in the same subtype of GABAergic cells and only exhibited different developmental expression patterns, we performed double *in-situ* hybridization (ISH) at P14, a developmental period when both genes are expressed high in vLGN. Indeed, our results showed that all cells that express *Lypd1* mRNA also generate *Ecel1* mRNA in vLGNe as well as vLGNi (Figure 2B, 2B’, 2B”), indicating that these markers correspond to the same GABAergic neuronal subtype.

**Figure 2.**
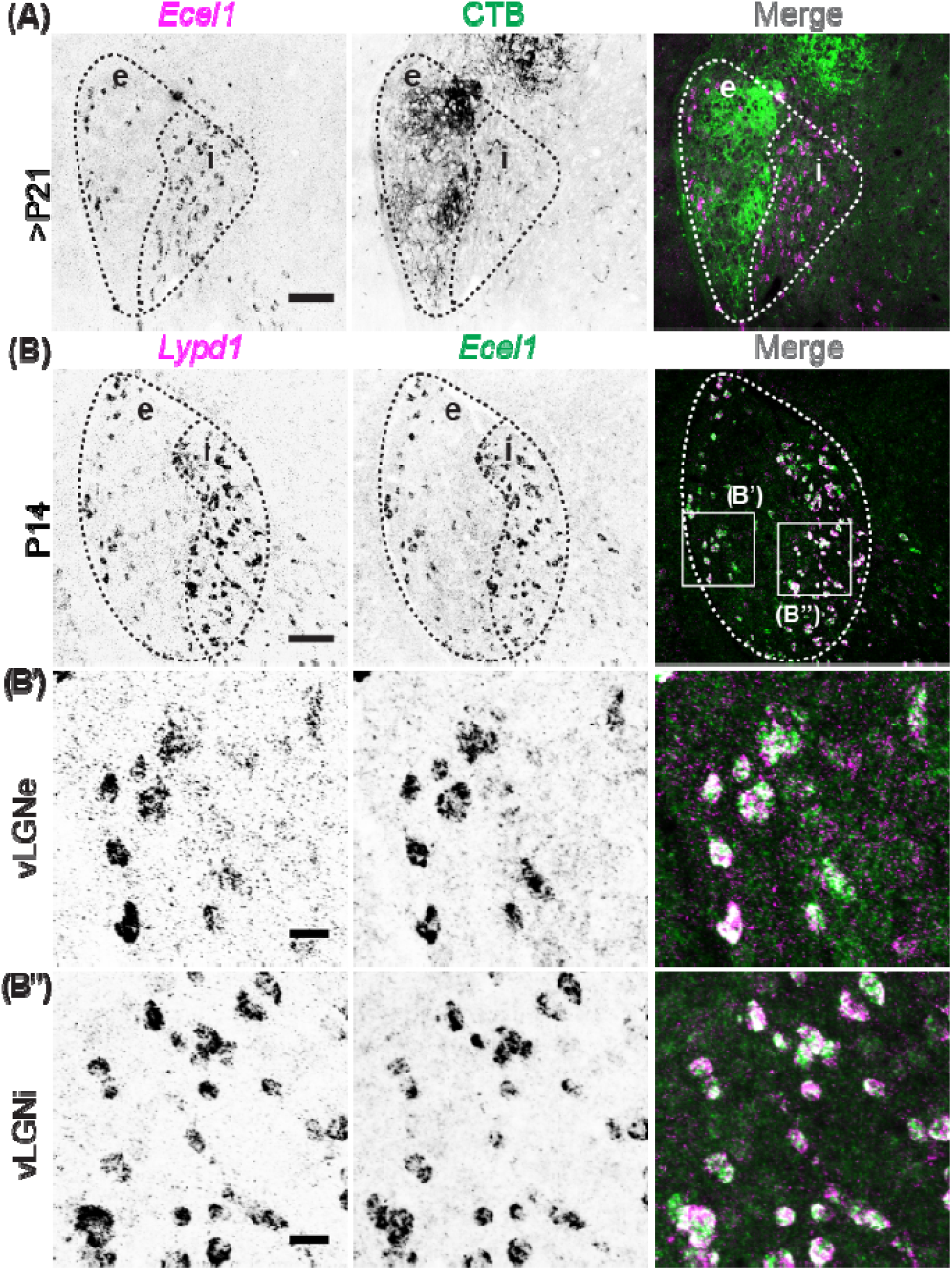
Molecular markers to identify the boundaries of vLGNe and vLGNi. (A) In-situ hybridization (ISH) with riboprobes against *Ecel1* following intraocular injection of fluorescently-conjugated CTB in >P21 WT vLGN. (B) Double ISH with riboprobes against *Lypd1* and *Ecel1* in P14 WT vLGN. Panels in B’ depict vLGNe. Panels in B” depict vLGNi. Scale bars in A,B: 100 μm and B’,B”: 20 μm.

To begin to elucidate whether the transcriptionally distinct laminae in vLGN were expressed during early development, we studied the development of *Ecel1*^*+*^/*Lypd1*^*+*^ and *Penk1*^*+*^ neurons. In adult mice, *Ecel1*^+^ cells occupy the lateral-most border of vLGNe and the entire vLGNi, while *Penk*^+^ cells occupy the medial-most border of vLGNe (Sabbagh et al., 2021). Interestingly, our double ISH results showed that the distinct spatial localization of *Lypd1*^+^ cells and *Penk*^+^ cells were present as early as P0 in the vLGN (Figure 3). These data suggest that the formation of cell type-specific laminae in vLGN occurs during embryonic development.

**Figure 3.**
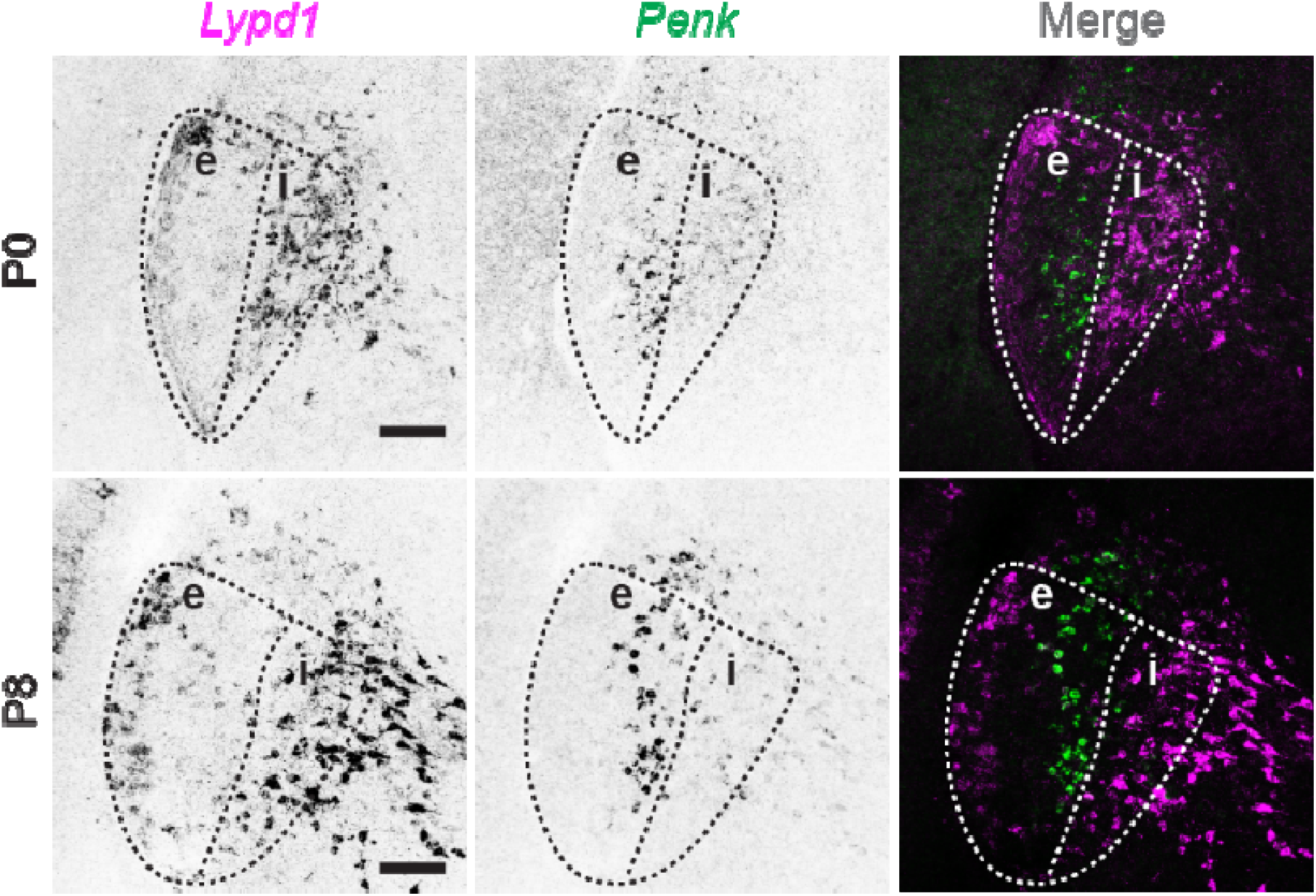
Subtype-specific sublaminae are specified in the neonatal vLGN. Double ISH with riboprobes against *Lypd1* and *Penk* at P0 and P8 in WT vLGN. Scale bars: 100 μm.

### Loss of retinal inputs disrupts vLGN cytoarchitecture lamination

The existence of distinct GABAergic laminae in the retinorecipient vLGNe, but not in vLGNi, suggests that retinal inputs might be involved in the lamination of vLGN. Previous studies have found that retinal inputs direct spacing and migration of GABAergic interneurons in the dLGN (Golding et al., 2014; Charalambakis et al., 2019; Somaiya et al., 2022). Therefore, we asked whether retinal inputs are critical for specifying the laminar organization of GABAergic neurons in vLGN by utilizing *Math5*^*-/-*^ mice that lack RGCs and their central projections in the brain (Brooks et al., 2013; Wang et al., 2001; Hammer et al., 2014; Seabrook et al., 2013). To test this, we used double ISH with riboprobes against *Penk* at all ages and either *Lypd1* at ages up to P5 or *Ecel1* for ages >P5. We observed that GABAergic laminae exist in *Math5*^-/-^ vLGN at birth, and the spatial localization of *Ecel1*^*+*^*/Lypd1*^*+*^ *and Penk*^*+*^ cells is very similar to WT controls (Figure 4A). Interestingly, the separation in the spatial distribution of *Ecel1*^*+*^/*Lypd1*^*+*^ and *Penk*^*+*^ neurons within vLGN decreased dramatically by eye-opening (Figure 4B-C). To quantitatively assess the differences in the laminar organization of GABAergic neurons between control and *Math5*^-/-^ mice, we performed line scan analysis in ImageJ to measure binarized signal corresponding to *Ecel1*^*+*^/*Lypd1*^*+*^ and *Penk*^*+*^ neurons along the medial-lateral axis of vLGN. Indeed, our data showed that while *Ecel1*^*+*^/*Lypd1*^*+*^ and *Penk*^*+*^ laminae exist at birth in *Math5*^-/-^ mice (Figure 4A’), these GABAergic subtypes begin to lose their spatial localization after the first week of postnatal development (Figure 4B’ and 4C’) and by eye-opening are occupying overlapping territory in vLGN (Figure 4D’).

**Figure 4.**
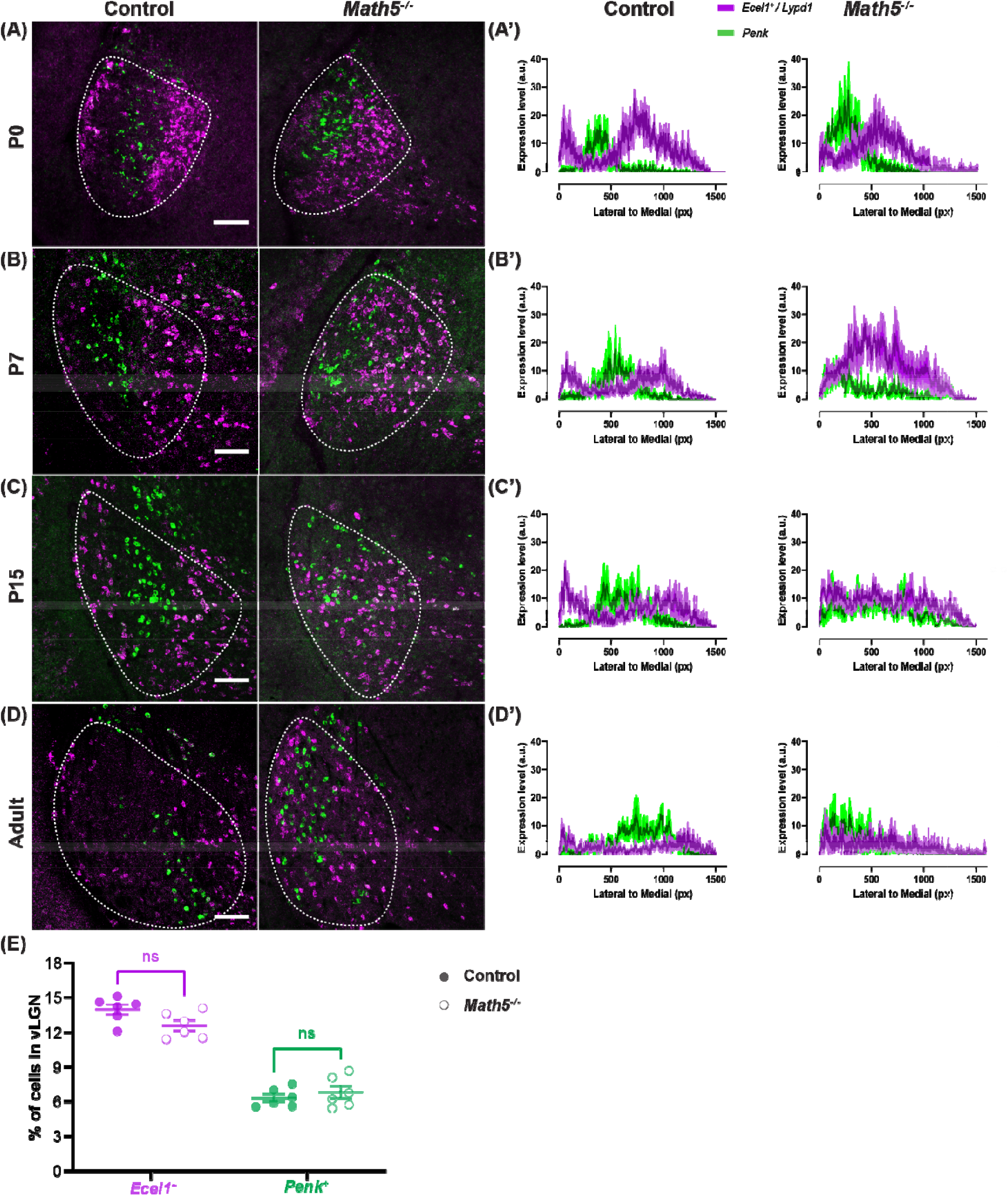
Loss of retinal inputs leads to progressive disruption of vLGN lamination. (A) Double ISH with riboprobes against *Lypd1* and *Penk* in P0 WT and *Math5*^-/-^ mice. (B, C, D) Double ISH with riboprobes against *Ecel1* and *Penk* in (B) P7, (C) P15, and (D) adult WT and *Math5*^-/-^ mice. (A’) Line scan analysis of spatial distribution of *Lypd1*^+^ and *Penk*^+^ cells. Arbitrary fluorescence units (a.u.) are plotted against distance from the lateral-most part of the vLGN to the most medial by pixel (px). Solid line represents mean and shaded area represents SEM (n=3). (B’, C’, D’) Line scan analysis of spatial distribution of *Ecel1*^+^ and *Penk*^+^ cells. Arbitrary fluorescence units (a.u.) are plotted against distance from the lateral-most part of the vLGN to the most medial by pixel (px). Solid line represents mean and shaded area represents SEM (n=3). (E) Quantification of the percentage of *Ecel1* and *Penk*-expressing cells in vLGN of P14 WT and *Math5*^-/-^ mice (n=6). Scale bars in A,B,C,D: 100 μm.

The disrupted organization could be a result of the lack of retinal input-dependent cues (molecules and/or intrinsic activity) that are required to maintain laminar arrangement or cues required for the survival of vLGNe neurons. To explore the latter possibility, we next determined whether there were changes in the percentage of different GABAergic subtypes in the absence of the retinal inputs. We assessed the percentage of cells rather than the cell density due to the significant difference in total vLGN area and cell count between control and *Math5*^-/-^ LGN (Charalambakis et al., 2019; Hammer et al., 2014; El-Danaf et al., 2015). By quantifying *Ecel1*^*+*^/*Lypd1*^*+*^ and *Penk*^*+*^ neurons, our results show no change in their percentage between control and *Math5*^-/-^ vLGN (Figure 4E). Taken together, these data suggest that retinal inputs are not impacting GABAergic cell proliferation, recruitment, or survival in vLGN or the formation of inhibitory laminae in vLGNe, but instead are essential for the maintenance of cell-type specific layers in vLGNe.

### Loss of *Fgf15* does not disrupt subtype-specific organization in vLGN

Recent studies revealed that RGCs induce the expressions of fibroblast growth factor 15 (FGF15), which is required for the recruitment of GABAergic interneurons into dLGN and vLGN (Su et al., 2020, Somaiya et al., 2022). Given that the absence of retinal inputs disrupts vLGN cytoarchitecture lamination, we wondered whether FGF15 was a retinal-induced signal required for the maintenance of cell-type specific layers in vLGNe. To test this, we performed double ISH with riboprobes generated against *Ecel1* and *Penk* mRNA in control and *Fgf15*^-/-^. We observed that neurons generating *Ecel1* and *Penk* mRNA retained their laminar identities in the *Fgf15*^-/-^ mice (Figure 5A-B), which was confirmed with the quantitative assessment via line scans (Figure 5A’ and 5B’). We also found no change in the percentage of *Ecel1*^+^ and *Penk*^+^ cells between control and *Fgf15*^-/-^ vLGN (Figure 5C). These findings indicate that the developmental influence of FGF15 is not critical for the recruitment and distribution of *Ecel1*^+^ and *Penk*^+^ cells in vLGN, nor is it necessary for the maintenance of these GABAergic laminae.

**Figure 5.**
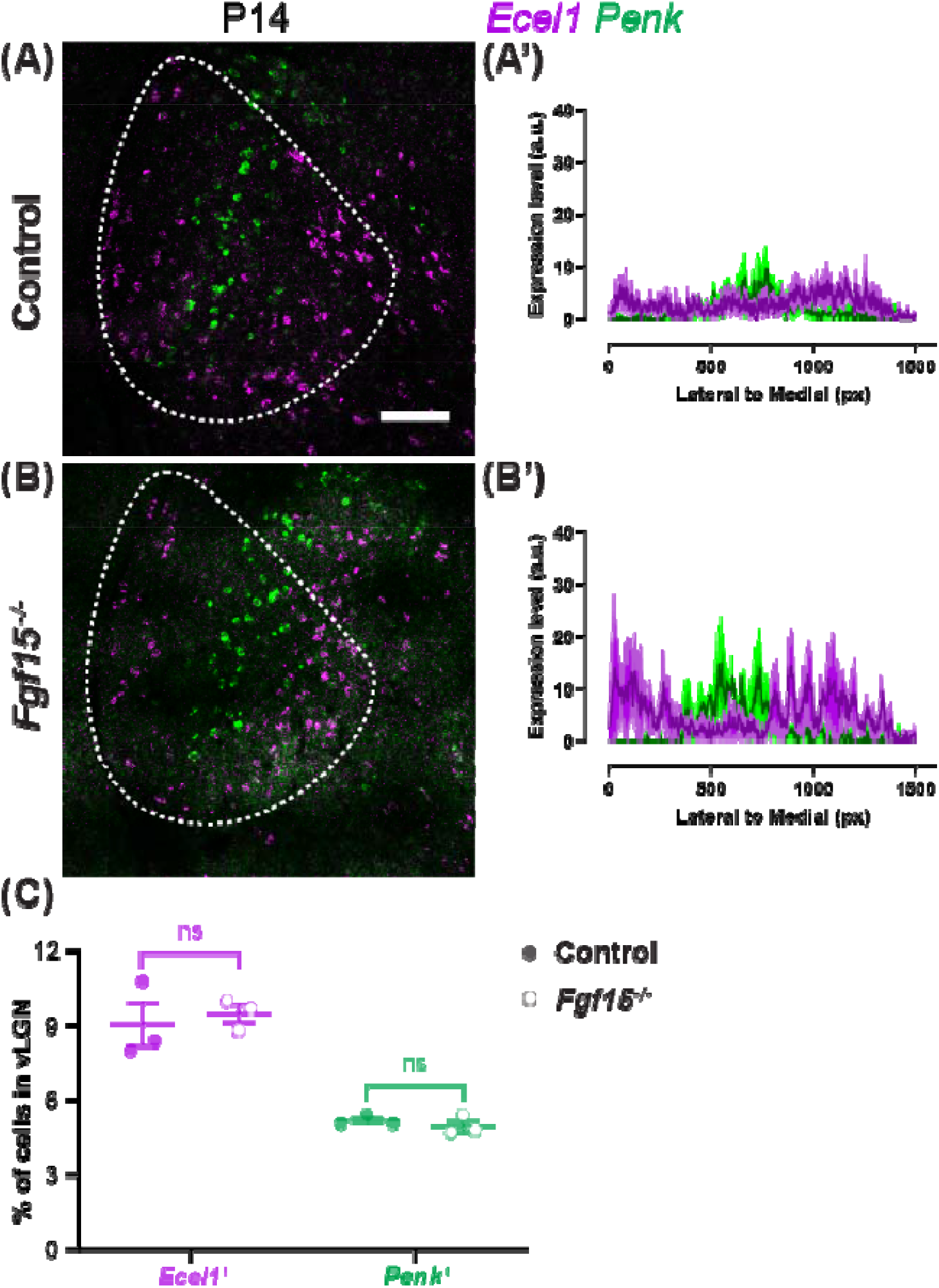
Loss of *Fgf15* does not disrupt subtype-specific organization in vLGN. (A, B) Double ISH with riboprobes against *Ecel1* and *Penk* in P14 WT and *Fgf15*^-/-^ mice. (A’, B’) Line scan analysis of spatial distribution of *Ecel1*^+^ and *Penk*^+^ cells. Arbitrary fluorescence units (a.u.) are plotted against distance from the lateral-most part of the vLGN to the most medial by pixel (px). Solid line represents mean and shaded area represents SEM (n=3). (C) Quantification of the percentage of *Ecel1* and *Penk*-expressing cells in vLGN of P14 WT and *Fgf15*^-/-^ mice (n=3). Scale bars in A,B: 100 μm.

## DISCUSSION

In this study, we characterized the molecular markers labeling distinct subtypes of GABAergic neurons in vLGN during development. Using *Ecel1*^+^ and *Penk*^+^ neurons, we found that cell type-specific lamination in vLGNe is determined during embryonic and/or neonatal development, before the migration of local interneurons, the arrival of other types of axons, eye-opening, or the emergence of experience-dependent visual activity. In mutant mice with no RGC axons, we found that retinal inputs are required for the maintenance of vLGN cytoarchitecture organization but are not required for the formation of these laminae. Lastly, we demonstrated that this redistribution of principal GABAergic cells is not due to the loss of FGF15, an RGC-induced, astrocyte-derived guidance cue essential for the migration of interneurons into rodent LGN (Su et al., 2020, Somaiya et al., 2022). Together, these data provide insights into the role of sensory inputs in the development of vLGN cytoarchitecture.

Previous studies have shown that retinal inputs are necessary for the migration of GABAergic interneurons into visual thalamus from tectal and thalamic progenitor zones (Seabrook et al., 2013; Somaiya et al., 2022). However, our results show that retinal inputs are not required for the migration of vLGN GABAergic cell types explored in this study, indicating that *Ecel1*^+^ and *Penk*^+^ cells in vLGN are not interneurons but instead are likely projection neurons. In contrast, previous studies have demonstrated that GFP-labelled interneurons in the GAD67-GFP mouse line do require retinal inputs to properly migrate into vLGN (Su et al., 2020). Taken together, these data begin to suggest that retinal inputs exert different influences on principal (projection) neurons than on local interneurons in vLGN. This is certainly the case in dLGN, although projection neurons in dLGN are excitatory and not GABAergic.

The presence of *Lypd1*^+^ and *Penk*^+^ cells in vLGN, both labeling distinct subpopulations of neurons as early as P0, indicated to us that these GABAergic neurons have already migrated into the vLGN with specified molecular identity. And, while *Lypd1*^+^ and *Penk*^+^ neurons did in fact stratify into laminae, the layers they formed appeared to be less refined in the early developing vLGN (Figure 4A-D). For instance, in the adult vLGN, *Penk*^+^ neurons formed a more sharpened sublamina that is more clearly delineated (Figure 4D), whereas at P0, *Penk*^+^ neurons are still layer-forming, but appear to be more widespread and occupy more of the lateromedial space of the vLGN than during adulthood (Figure 4A). These differences suggest that this laminar organization continues to refine after birth. Since our results indicate that retinal inputs are not required for their formation but are necessary, retinal input-derived cues might be important for the refinement of GABAergic laminar organization in vLGN.

While previous studies have determined that astrocyte-derived FGF15 is essential for the recruitment of GABAergic interneurons, the loss of functional FGF15 did not reproduce the defects in subtype-specific lamination associated with the loss of retinal input. Thus, proper maintenance of vLGN lamination in the presence of retinal inputs could be due to other developmentally regulated factors released by RGC axons or retinal activity. Indeed, RGCs in the retina undergo significant transcriptomic changes during development, suggesting that molecules expressed after eye-opening could be potentially important for vLGN lamination (Shekhar et al., 2022). Another possibility is that retinal input is necessary for the proper development of vLGN extracellular matrix (ECM) rather than principal neuron lamination. To assess if disruption of retinal inputs impairs ECM formation and maintenance, which subsequently impairs the location of principal neurons in vLGN, it would be beneficial to evaluate chondroitin sulfate proteoglycans that are known to be enriched in vLGN, such as phosphacan (Brooks et al., 2013).

Overall, our study suggests two distinct developmental processes involved in specifying vLGN laminae: one to form a layer, the other to refine and sharpen it. Thus, the cells of vLGNe and vLGNi are genetically hardwired to be retinorecipient or non-retinorecipient, respectively, as opposed to assuming cellular phenotypes because of retinal input. Our data, when taken together, suggest the possibility that the functional organization of non-image-forming information from the retina to vLGN is extracted from the segregation of transcriptionally distinct retinorecipient cells. We view these results as a framework for further dissecting the structure, circuitry, and functions of the vLGN at a cell type-specific level. How the heterogeneity and organization contribute to the yet-to-be-determined functions of the vLGN remains to be defined.

## Acknowledgments

This work was supported by the National Institutes of Health grant EY033528 (M.A.F.), EY021222 (M.A.F.), NS113459/ K00NS113459 (U.S.), and EY035570 (K.S.). We are grateful to the members of the M.A.F. lab for scientific discussion and comments on the manuscript. We thank Dr. Steven W. Wang for providing *Math5*^*-/-*^ mice.

## Notes

### Competing Interest Statement

The authors have declared no competing interest.

